# Bayesian Integration in Sense of Agency: Understanding Self-attribution and Individual Differences

**DOI:** 10.1101/2025.02.19.639190

**Authors:** Acer Chan-Yu Chang, Wen Wen

## Abstract

This study developed a Bayesian integration model to explore individual differences in the sense of agency, particularly when multiple sensory cues influence action perception. Behavioral results showed that individuals consistently integrate sensory cues in agency judgments, though cue weighting varies significantly across individuals. The estimated parameters of our model successfully captured these differences, reflecting individual sensitivities and criteria for agency. Higher sensitivity to sensory inputs was associated with lower variance in the likelihood distribution under the assumption of control. Our model provides critical insights into the mechanisms of the sense of agency and its variability, highlighting the model’s potential for understanding both typical and disordered agency experiences.

## Introduction

Understanding the cause of sensory changes is crucial for an individual’s survival, particularly when differentiating between external events and self-generated actions. For example, visual shifts on the retina could result from either an external object’s movement or the rotation of one’s eyes. Through causal inference, individuals can better comprehend their interactions with the environment and their level of control over it. This awareness contributes to the sense of agency, which is the subjective experience of being in control of one’s actions(Gallagher, 2000).

In reality, an action can generate multiple forms of sensory feedback that span across space, time, and levels of abstraction, especially when the action is goal oriented. For instance, throwing a ball to hit a target involves both proximal (body-related) and distal (goal-related) feedback. We infer our degree of control over the action and form a unified sense of agency by integrating this diverse feedback throughout the entire action(Charalampaki et al., 2023). However, the exact process by which we integrate this feedback remains unknown. More importantly, individual differences in weighting various types of feedback could be significant, but the origin of such differences remains unclear.

Traditional theories of the sense of agency focus on proximal feedback. The comparator model, for instance, suggests that the sense of agency results from comparing predicted sensory feedback, based on an efference copy of the motor command, with the actual sensory feedback received(Blakemore et al., 2002; Frith et al., 2000; Wolpert and Flanagan, 2001). However, subsequent studies revealed that both proximal feedback (computed from motor commands) and distal feedback (such as intentions and goals) influence the sense of agency(Metcalfe et al., 2013; Pacherie, 2008; Vinding et al., 2013; Wen et al., 2015). These studies highlight that the sense of agency is not simply a matter of comparing predicted and actual bodily feedback but rather involves integrating multiple sources of information across various domains. Naturally, traditional theories are incapable of capturing how the sense of agency emerges from the integration of diverse forms of feedback across the entire action and how each individual integrates evidence differently.

Recent Bayesian models provide a promising framework to address these limitations. For examples, Bayesian cue integration models suggest that the sense of agency results from integrating multiple sensory cues, weighted by reliability(Moore and Fletcher, 2012; Synofzik et al., 2009). Legaspi and Toyoizumi present a model explaining intentional binding and precision-dependent causal agency(Legaspi and Toyoizumi, 2019). Roth et al. further highlight the role of Bayesian causal inference in understanding impaired temporal perception in schizophrenia(Roth et al., 2023). This raises the question of whether a similar principle also applies to the integration of different types of feedback, from proximal to distal.

In this study, we hypothesize that the unified sense of agency throughout an entire action is derived from inferring the cause of external events. This inference integrates multiple forms of feedback using Bayesian principles. In Bayesian inference involving evidence from multiple modalities, the importance (weighting) is determined by the precision (reliability) of the evidence. Evidence from a modality with higher precision gains more weight in the computation of the posterior belief (Ernst and Banks, 2002; Ernst and Bülthoff, 2004). We further predict that the integration of proximal and distal feedback follows Bayesian principles, with individuals assigning different weights to each type of feedback based on their reliability, ultimately shaping their sense of agency.

In addition, a Bayesian approach can potentially account for individual differences in the sense of agency, which have been largely overlooked. Empirical research has revealed significant individual differences in agency judgment. For example, sensitivity to detecting control varies widely in healthy populations(Wen et al., 2021). Additionally, when sensory feedback of an action is slightly delayed, the sense of agency differs significantly depending on individuals’ prior experience with delayed feedback(Wen, 2019). Such individual differences in both sensitivity and criteria for the sense of agency are noteworthy for understanding and predicting human behavior(Wen et al., 2024). The individual differences may arise from assigning different weights to various types of feedback, based on an individual’s unique estimation of the precision of different modalities.

In the current study, we designed a shooting task to evaluate how participants integrate proximal and distal cues in judging their sense of agency. In Experiment 1, participants used a joystick to launch a dot toward a moving target arc (Figure 1A). We manipulated two types of errors: the proximal error (angle between joystick direction and launch direction, varying from 0-40 degrees) and the distal error (angle between launch direction and target center, varying from 0-20 degrees) (Figure 1B). The dot took 500 ms to reach the target arc, during which the program adjusted the target’s position to create the designated distal error. After each trial, participants rated their sense of agency over the dot’s trajectory on a 0-1 scale. Furthermore, Experiment 2 was conducted for two main purposes. First, it aimed to replicate the findings of Experiment 1. Second, it included a static shooting task (Figure 1C) after the main task to measure the empirical motor control performance of each individual, allowing us to estimate the empirical reliability (variance and precision) of their proximal feedback. With this data, we could determine if participants integrated proximal and distal cues based on their proximal precision, an important feature of Bayesian cue integration. We also proposed a parsimonious Bayesian computational model to simulate observed behavioral patterns in the agency task. This model can explain substantial individual differences by assigning different weights to proximal and distal cues.

**Figure 1.**
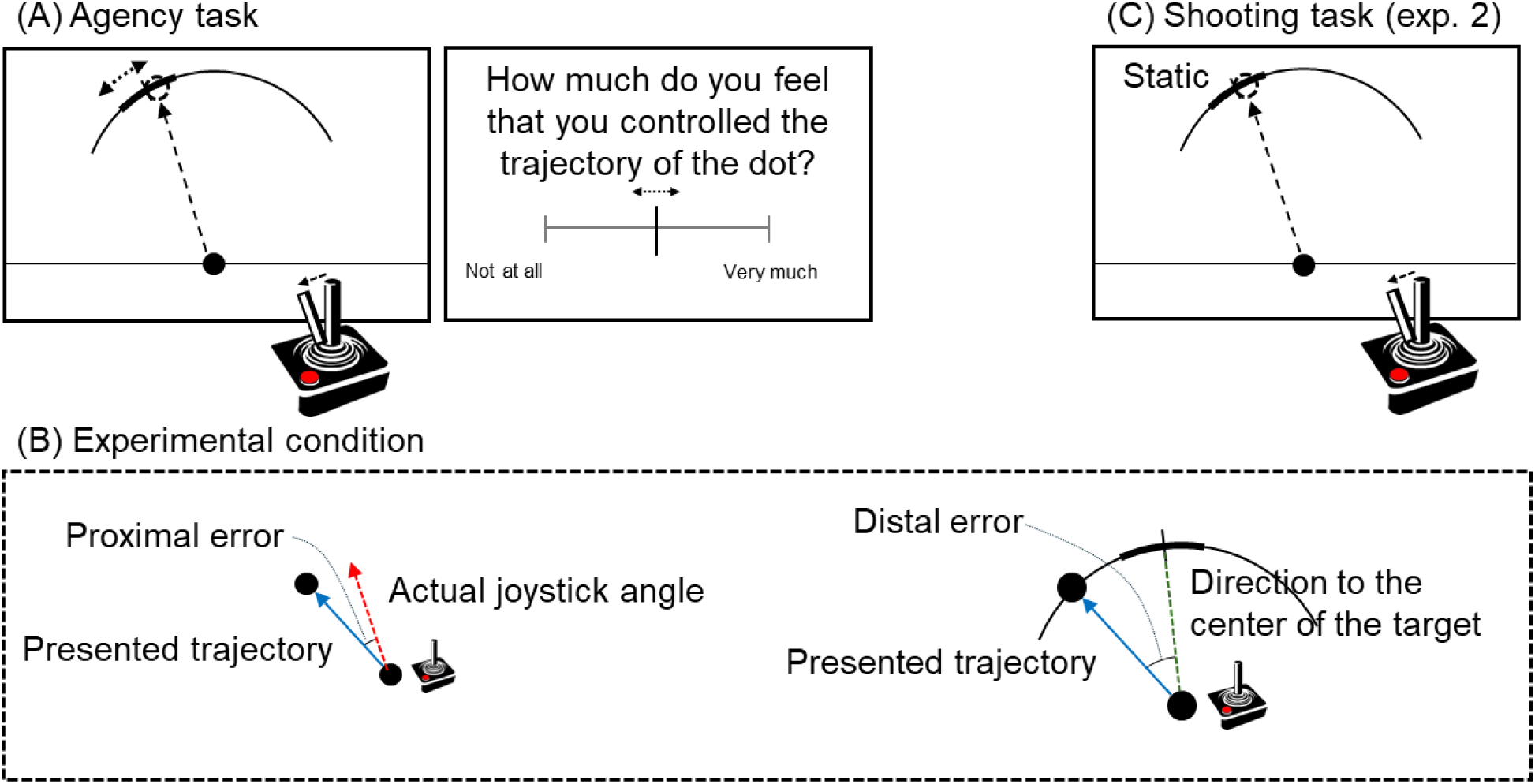
Task design and experimental setup. (A) Agency task: Participants used a joystick to launch a dot toward a moving target arc, followed by rating their sense of control. (B) Error measurements: Two types of errors were manipulated - proximal error (difference between joystick and launch direction) and distal error (difference between launch direction and target center). (C) Static shooting task used in Experiment 2 to assess individual motor performances

### Bayesian cue integration model

Assuming that sensory inputs can be caused by both the self and other (i.e., the environment), when giving a sensory input (D), the probability of self (H = self) can be calculated from:

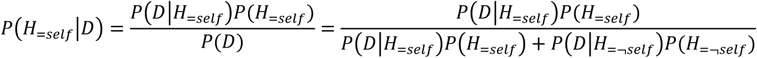

When there are two independent cues (D1 and D2), each can be caused by the self and other, the probability of self (*P*(*H*_=*self*_)) when given the two cues can be calculated from:

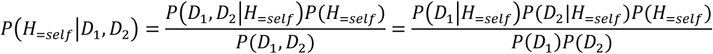

This can be further expanded to:

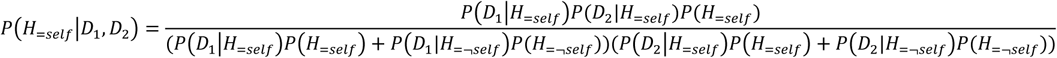

The sum of *P*(*H*_=*self*_) and *P*(*H*_=¬*self*_) should be 1. For simplicity, we can use a Guassian distribution for sensory input caused by the self *P*(*D*|*H*_=*self*_). Additionally, if there is no specific assumption for the environment, we can assume a uniform distribution for *P*(*D*|*H*_=¬*self*_). With this model, we propose that the variance (σ²) of the likelihood distribution for a sensory cue determines its weighting in agency judgment. Figure 2A and 2B illustrate examples of self-attribution with one and two cues.

**Figure 2.**
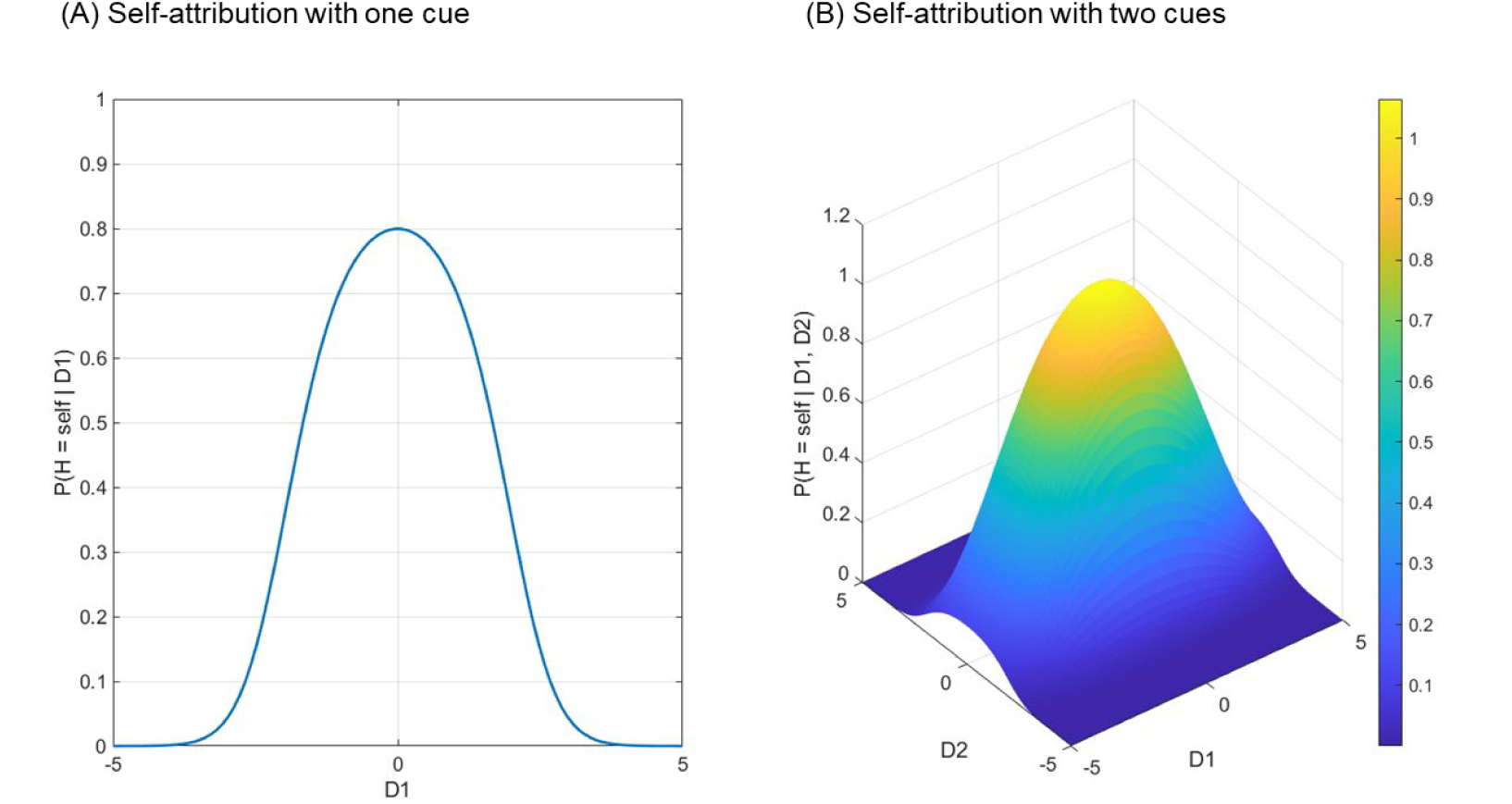
Simulation of the model. Panel (A) shows the example of self-attribution with one cue when assuming a uniform distribution for the others. Panel (B) shows the example of self-attribution with two cues.

## Results

### Behavioral results

Figure 3 shows individual agency ratings for both experiments (n = 28 for Experiment 1 and n = 30 for Experiment 2). In general, the results indicate that participants considered both proximal and distal errors in their agency ratings. For Experiment 1, the 5 × 5 (proximal error × distal error) repeated measures ANOVA showed that the main effect of proximal error (*F*(1.371, 37.009) = 168.752, *p* < .001, partial η^2^ = .862), the main effect of distal error (*F*(1.603, 43.288) = 19.431, *p* < .001, partial η^2^ = .418), and the interaction between the two factors (*F*(16, 432) = 1.678, *p* = .048, partial η^2^ = .059) were all significant. The Greenhouse-Geisser correction was applied when the sphericity assumption was violated. For Experiment 2, the same repeated measures ANOVA showed that both the main effect of proximal error (*F*(1.224, 35.495) = 41.835, *p* < .001, partial η^2^ = .591) and the main effect of distal error (*F*(1.151, 33.393) = 52.021, *p* < .001, partial η^2^ = .642) were significant, but the interaction between the two factors was not (*F*(9.217, 267.302) = 1.741, *p* = .078, partial η^2^ = .057). Although participants were explicitly asked to base their sense of agency ratings on the dot’s trajectory while disregarding target-hitting performance, the distal error still influenced many participants. Importantly, there were notable individual differences in how they weighted each cue type and determined the boundary of their sense of agency. Figure 4 presents data from two representative participants (see Supplementary Material S1 for the density distribution of all participants). Some participants exhibited shifts in the peaks of the probability density distributions when the proximal cue was altered, while others showed such shifts when the distal cue was altered. This implies that some participants are more sensitive to the proximal cue while others are more sensitive to the distal cue.

**Figure 3.**
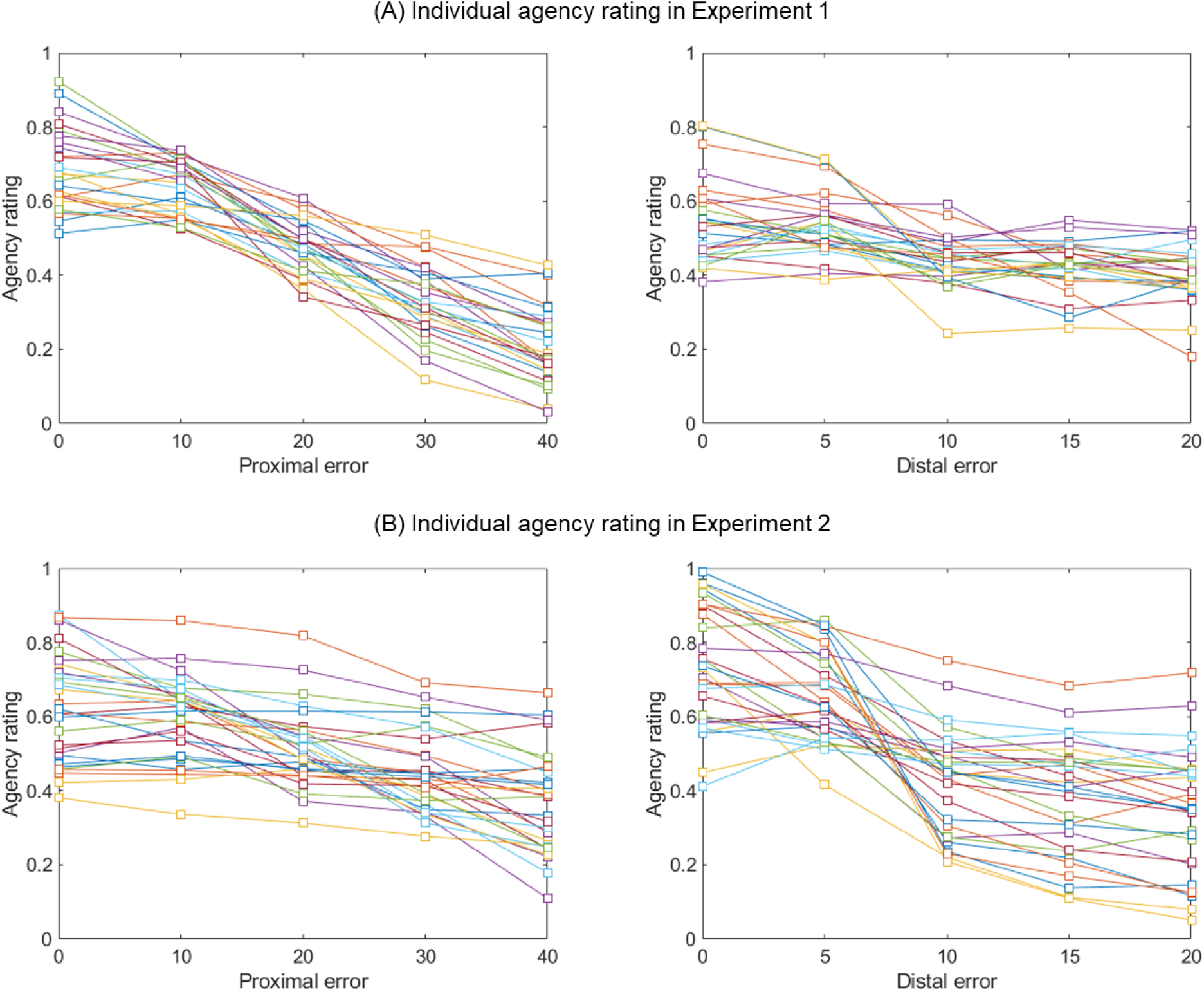
Individual agency ratings in Experiment 1 and 2. In both experiments, participants rated their sense of agency over the trajectory of the dot depending on both the proximal error and the distal error. Colored lines represent individual participants.

**Figure 4.**
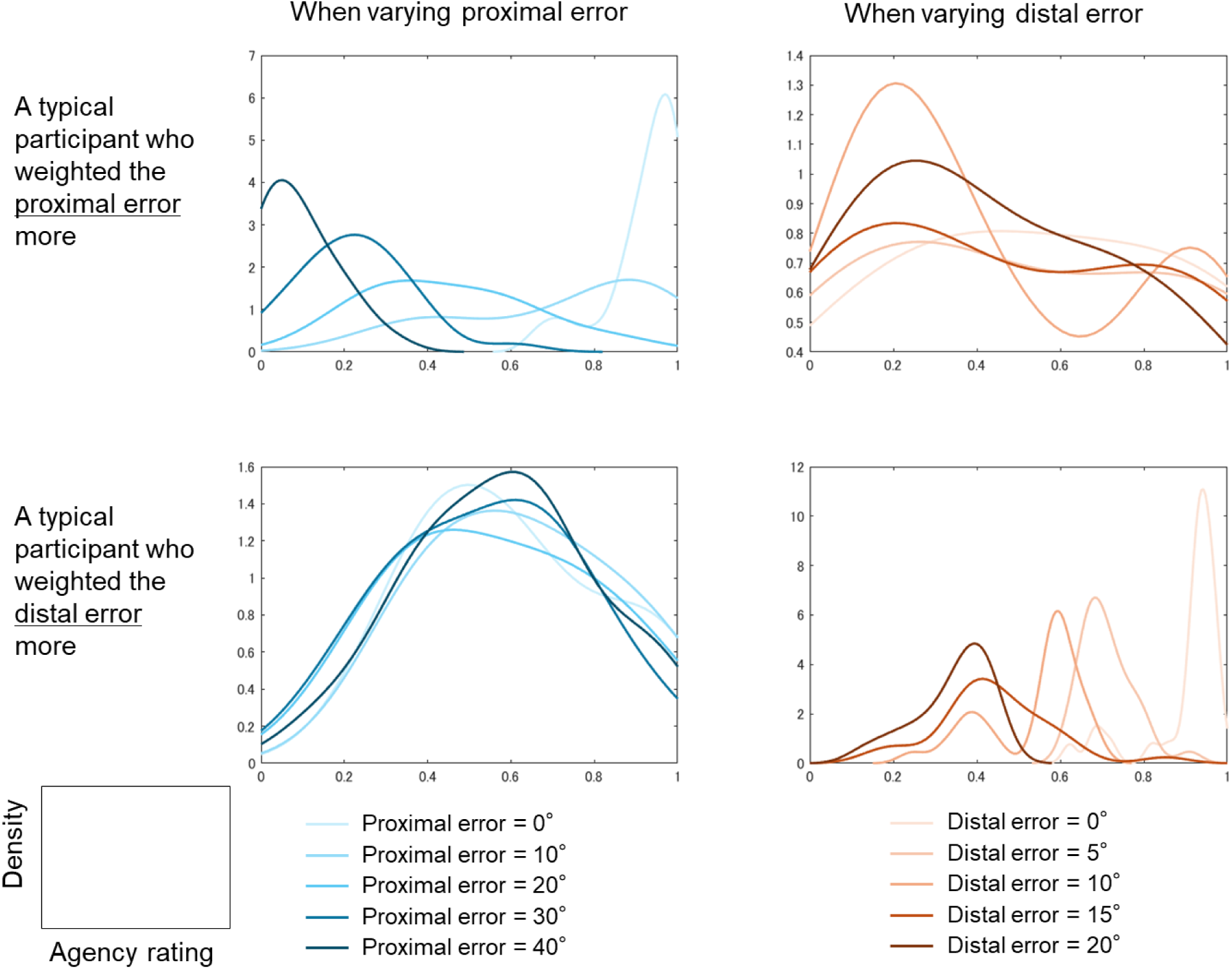
Probability density distribution of agency judgment for two participants who weighted the two types of cues differently. To enhance visualization of these distributions, kernel density estimation was applied.

Next, to assess the extent to which individuals rely on a cue to rate their sense of agency, we calculated the correlation between agency ratings and each type of error. This correlation reflects how closely one’s agency judgment relies on each cue. Figure 5A shows the distribution of each type of correlation and the correlation between the two types. A significant finding is that the correlations of the two cues with agency judgment are strongly negatively correlated with each other (Figure 5B: *r* = -0.84 and -0.94 for Experiments 1 and 2, respectively). These strong correlations highlight a characteristic of Bayesian integration.

**Figure 5.**
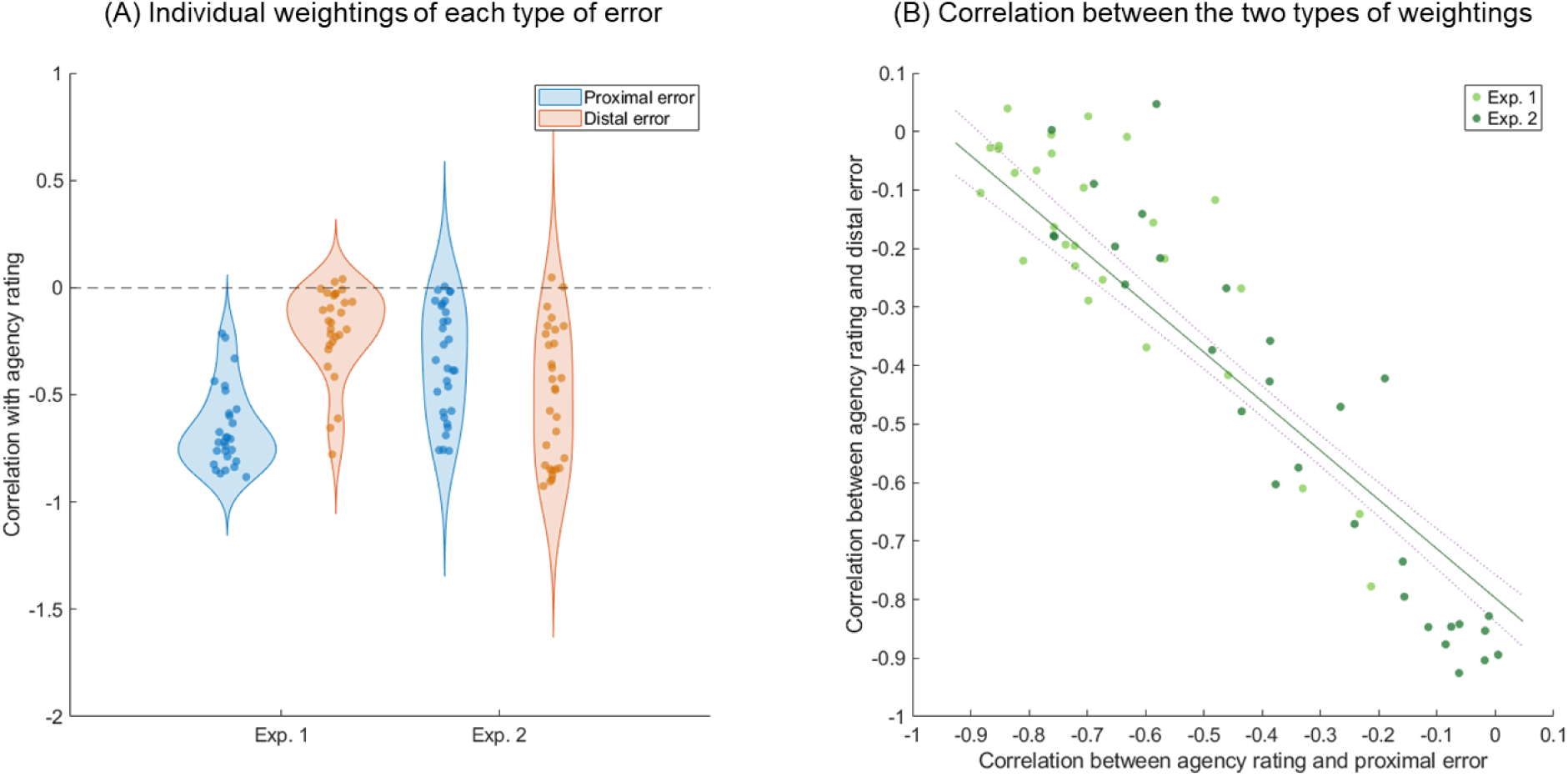
Correlations of agency judgment with the two types of cues. Panel (A) shows the distribution of individual correlations for proximal and distal errors with agency judgment. Panel (B) illustrates the strong negative correlation between the two correlations in both experiments.

Lastly, Experiment 2 additionally measured individual motor control performance. Participants aimed at a static target 40 times, with full control over the joystick and a goal of consistently aiming at the target’s center. In this case, the variance in performance error may approximately reflect the likelihood distribution of proximal errors when one has full control. We fitted a Gaussian distribution to their performance and obtained the standard deviation (σ) for each participant (Figure 6A). This parameter reflects the distribution of possible proximal errors when participants have full control. The fitted parameter σ was significantly correlated with the correlation between agency ratings and proximal errors (*r* = .397, *p* = .030; Figure 6B), indicating that the sensitivity to a sensory input in agency judgment is linked to the likelihood distribution of that cue.

**Figure 6.**
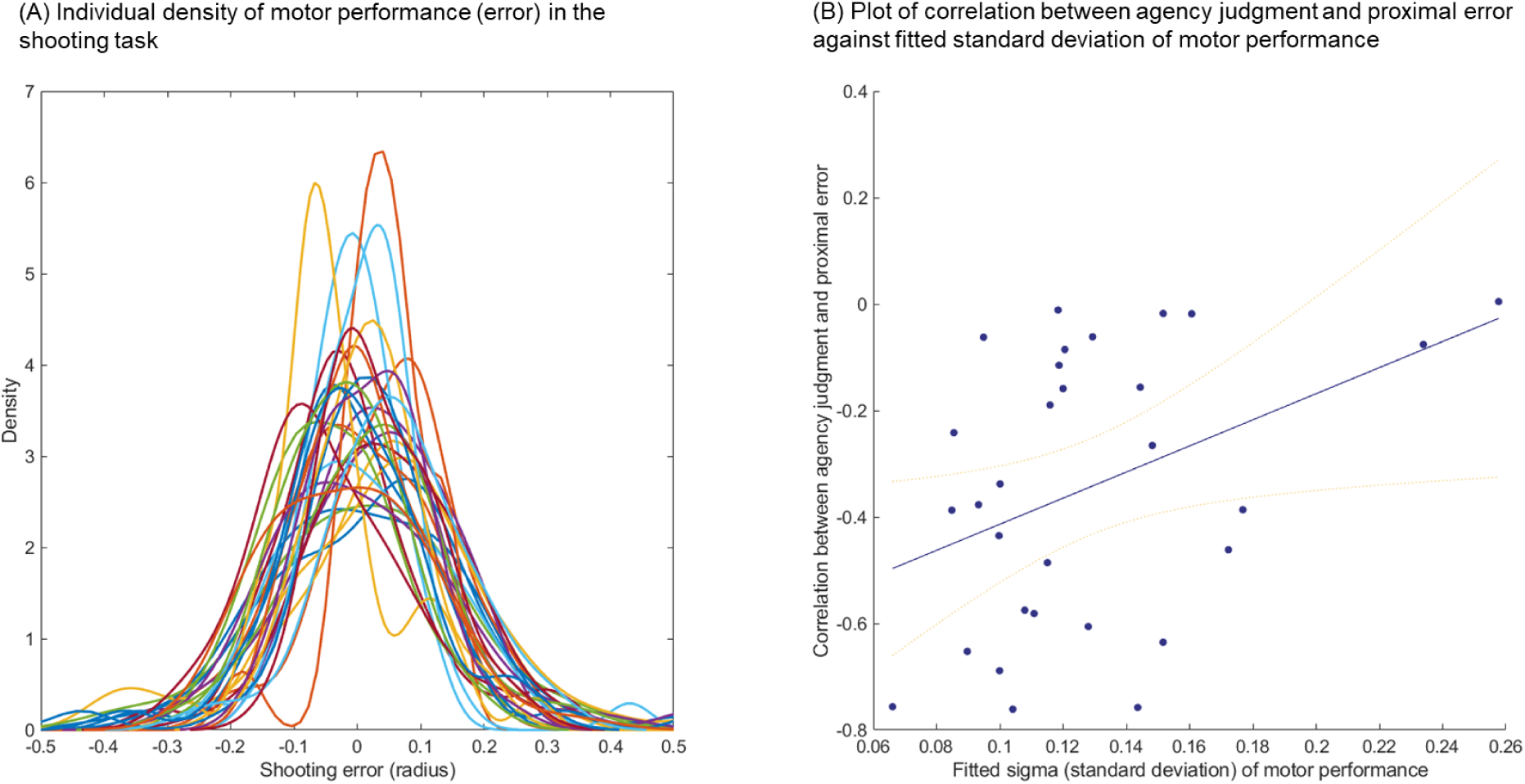
Individual motor performance in the shooting task (A) and the plot of motor performance (fitted parameter σ) and weighting of proximal error in agency judgment (B).

### Parameter Estimation and Distribution of Individual Difference

Figure 7A shows a heatmap of predicted agency judgments against participants’ actual agency judgments. The predicted agency judgments demonstrate good model fit to the actual agency ratings (mean RSME = 0.18, *SD* = 0.04; mean R-squared = 0.52, *SD* = 0.16), indicating that the proposed model effectively captured the way people make agency judgments. Figures 7B and 7C further display the densities of σ1, σ2 (standard deviations of *P*(*D*_1_|*H*_=*self*_) and *P*(*D*_2_|*H*_=*self*_), respectively), and prior belief (*P*(*H*_=*self*_)), illustrating the individual differences in these parameters within our sample.

**Figure 7.**
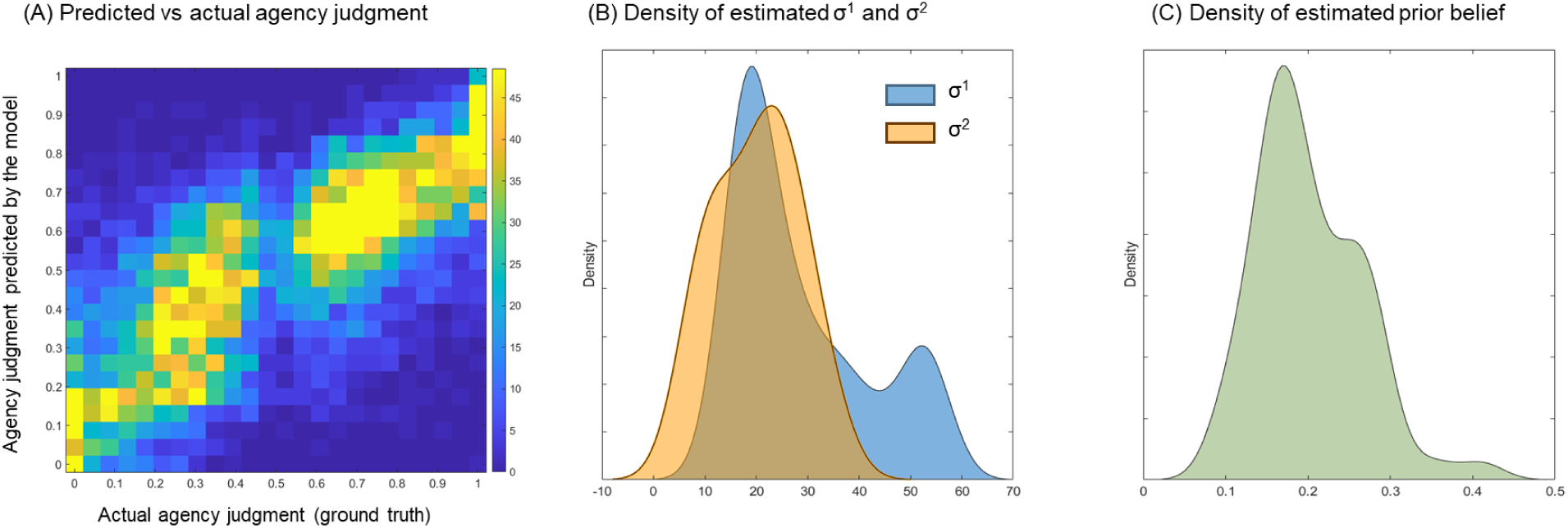
Results of Parameter Estimation and Model Prediction. Panel (A) shows a heatmap of predicted agency against actual agency judgments. Colors represent density of data. The high correlation between the density of them indicates a good fit of our model to the actual data. Panel (B) presents the density of the estimated standard deviations (σ1 and σ2) of the likelihood distributions for proximal and distal errors across individuals. Panel (C) displays the density of estimated prior beliefs across individuals. To enhance visualization of these distributions, kernel density estimation was applied for panels (B) and (C).

### Connect the signal detection theory of the sense of agency and the Bayesian cue integration

Wen et al. proposed that when examining individual differences in the sense of agency, one should evaluate sensitivity and criterion of their agency judgements based on signal detection theory(Wen et al., 2024). Here, we emphasize that the Bayesian model aligns with this approach. The values of σ1 and σ2 are linked to the sensitivity of the sense of agency, indicating how well one can distinguish between control and non-control based on sensory input. On the other hand, the value of prior belief is linked to the criterion for the sense of agency, indicating how likely one is to attribute sensory inputs to themselves.

Individual sensitivity to each type of sensory cue can be calculated by fitting a sigmoid function (see Method) using the agency rating against the value of the sensory cues, and individual criterion of sense of agency can be measured by the average agency rating. To evaluate whether the estimated parameters captured individual sensitivity and the criterion of the sense of agency, we plotted the estimated parameters of the Bayesian cue integration model against the indices of sensitivity and criterion of sense of agency acquired from the behavioral results. The plots showed that the estimated σ1 was significantly correlated with sensitivity to the proximal error, and the estimated σ2 was significantly correlated with the sensitivity to the distal error (*r* = .927 and .932, *p*s < .001, Figure 8A and 8B, respectively). Furthermore, the estimated prior belief was significantly correlated with the mean individual agency ratings (*r* = .719, *p* < .001, Figure 8C). Taken together, these validations indicate that our model effectively captures individual differences in both the sensitivity and criterion of the sense of agency.

**Figure 8.**
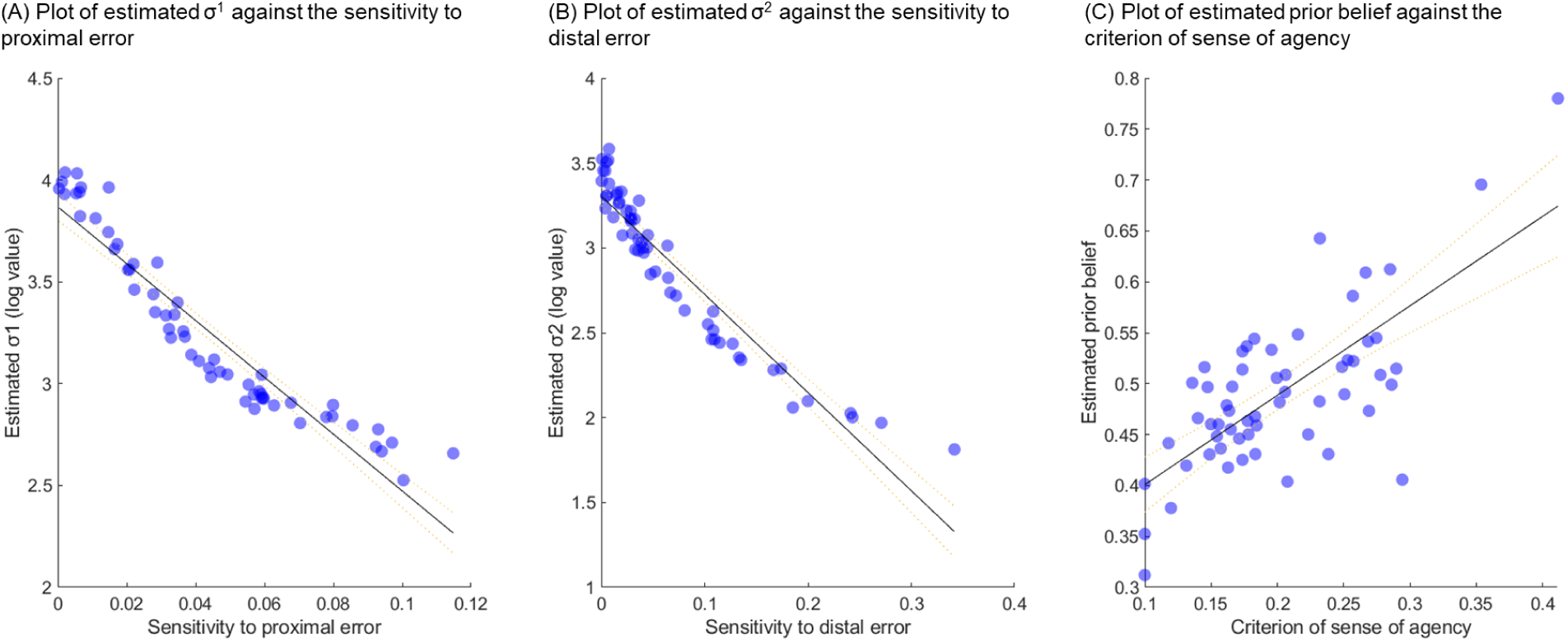
Plot of estimated parameters with measures of individual sensitivities and criteria of sense of agency. The *x*-values and y-values in panels (A) and (B) are the absolute values of the slopes and log values of estimated σ1 and σ2, respectively, to ensure positive values for better sensitivity and a better visualization of the linear relationship. Smaller estimated σ values are linked to better sensitivity to the corresponded cue, and higher estimated prior beliefs are linked to stronger sense of agency.

### Model simulation of abnormal sense of agency

Additionally, in order to examine whether the proposed Bayesian Integration model can represent abnormal patterns of agency judgment in psychosis such as Schizophrenia(Leptourgos and Corlett, 2020; Tan et al., 2023), we conducted a model simulation with reduced sensorimotor processing (i.e., a large σ for the sensorimotor cue) and enhanced retrospective processing (i.e., a small σ for the retrospective cue) while varying the sensorimotor input and prior belief. The simulated agent shows a density of self-attribution with two peaks across all three conditions of prior belief. Sensitivity to the sensorimotor cue (i.e., D1) in self-attribution is particularly low when the prior belief is strong. The simulated agent shows density peaks for both a lack of agency and a heightened sense of agency, relying solely on retrospective cues without adequate sensorimotor evidence. This pattern is especially strong when holding strong beliefs. This simulated agent aligns with the typical symptoms of abnormal sense of agency in psychosis (see Discussion).

## Discussion

The present study aimed to use a Bayesian integration model to understand individual differences in the sense of agency when multiple sensory cues are relevant to the action. We showed that people integrate multiple sensory cues for agency judgment in a consistent way, while the weightings of these cues vary greatly across individuals. We demonstrated that the estimated parameters of the Bayesian integration model effectively capture individual differences in both the sensitivity to each type of cue and the criterion for the sense of agency. Specifically, the smaller the variance of an individual’s likelihood distribution of a sensory input under the assumption of control, the more sensitive that individual is to that type of sensory input in agency judgment. When there are multiple cues, the probability of self is calculated based on Bayesian principles. This concept is straightforward and has been discussed in previous studies, such as the cue integration theory and recent review papers(Moore and Fletcher, 2012; Synofzik et al., 2009; Wen and Imamizu, 2022). However, the present study is the first to use computational modeling and parameter estimation to formalize and validate this theory. Our results provide important insights into how the sense of agency arises and where the individual differences underlying this process originate.

The concept of the sense of agency originates from philosophy(Gallagher, 2000) and has been widely studied in psychology and neuroscience since then because of its close link to self-consciousness. Many studies have revealed its impact on perception and behavior(Moore, 2016; Wen and Imamizu, 2022) and have discussed its neural and theoretical mechanisms(David et al., 2008; Haggard, 2017). In the 1980s, Vallacher and Wegner suggested that personal agency beliefs could affect how people use different identities ranging from low-level (e.g., identities that specify how the action is performed) to high-level (identities that signify the intention and goal of the action) for action identification(Vallacher and Wegner, 1989). Our behavioral results also revealed large individual differences in how people use goal-achievement information (i.e., distal errors) and sensorimotor outcomes (i.e., proximal errors) in their agency judgments. Such individual differences in processing sensory input could arise from high-level personality traits or beliefs(Rollwage et al., 2018; Vallacher and Wegner, 1989) but are more likely to stem from individual experiences with sensory input when they act. Studies in human-automation interaction suggest that more skillful operators often benefit less from automation or may even be disturbed by it(Wen et al., 2019; Wright et al., 2018). This is likely because more skillful operators have less variance in the likelihood distribution for the sensory input of their actions and, therefore, are more likely to notice discrepancies between the sensory input manipulated by automation and the possible sensory input when they have full control. Our behavioral results from Experiment 2 are consistent with this hypothesis, showing that people whose shooting performance was less variable were more sensitive to proximal errors in their agency judgments. This means that people must have access to their internal models of sensory cues. Such internal models can be learned in our daily life and can be generalized to similar situations via metacognition.

Furthermore, the assumption of other agents in the environment is important in our model. This factor has not been considered by previous models of the sense of agency, though it has been discussed in theories of the sense of agency. Specifically, the retrospective theory suggests that exclusivity (i.e., the absence of alternative causes for outcomes) is one of three necessary principles for the sense of agency (the other two being consistency and priority)(Wegner, 2003; Wegner et al., 2004). In our model, both the prior belief in other agents (*P*(*H*_=¬*self*_)) and the likelihood distribution of sensory input being caused by other agents (*P*(*D*|*H*_=¬*self*_)) affect self-attribution. In our task, the estimated prior belief of self was biased toward the environment (Figure 7C), likely because we frequently disturbed participants’ control by inducing proximal and distal errors. This prior belief can be updated if the sensory input is not random and can be learned. Although we did not include an algorithm for prior updating in the parameter estimation, it could be implemented depending on the control circumstances to improve model fit. Moreover, we assumed a uniform distribution for other agents, as participants in our task were unlikely to have specific assumptions about the ‘intentions’ of the computer algorithm. However, in real-world situations, assumptions about other agents can be more complex, depending on the alternative agents present in the environment. Our model can accommodate such circumstances by specifying the likelihood distribution for other agents.

The discussion of individual differences in the sense of agency is important not only for predicting human behaviors but also for examining agency and psychosis. Here, we showed that our Bayesian integration model is useful for understanding the phenomenon of sense of agency including in psychosis. Psychopathological symptoms in patients with schizophrenia suggest that their ability to distinguish between self and other is impaired(Schneider, 1959; Stanghellini and Monti, 1993). Specifically, patients with schizophrenia often misattribute self-generated sensory input, such as their own inner voice and eye movements, to external causes(Lindner et al., 2005; McGuire et al., 1995). Conversely, in some circumstances, patients also make excessive agency judgments compared to healthy controls(Hur et al., 2014; Koreki et al., 2015; Leptourgos and Corlett, 2020; Mariano et al., 2024; Renes et al., 2013; Rossetti et al., 2024). This abnormal pattern of agency judgments can be simulated with our model by assuming a weakened sensorimotor process(Dakin et al., 2005; Krugwasser et al., 2022; Leube et al., 2010; Oi et al., 2024; Posada et al., 2007; Powers et al., 2016; Uhlmann et al., 2021) and an enhanced retrospective component(Hauser et al., 2011; Metcalfe et al., 2012; Voss et al., 2010). Our model simulation successfully showed that the combination of reduced sensitivity in sensorimotor processing and enhanced retrospective processing can lead to abnormal judgments of agency (Figure 9). Specifically, the simulated agent may alternate between a lack of agency and a heightened sense of agency, relying solely on retrospective cues without adequate sensorimotor evidence, especially when holding strong beliefs. This simulated pattern aligns with behaviors observed in schizophrenia, such as jumping to conclusions(Rubio et al., 2011).

**Figure 9.**
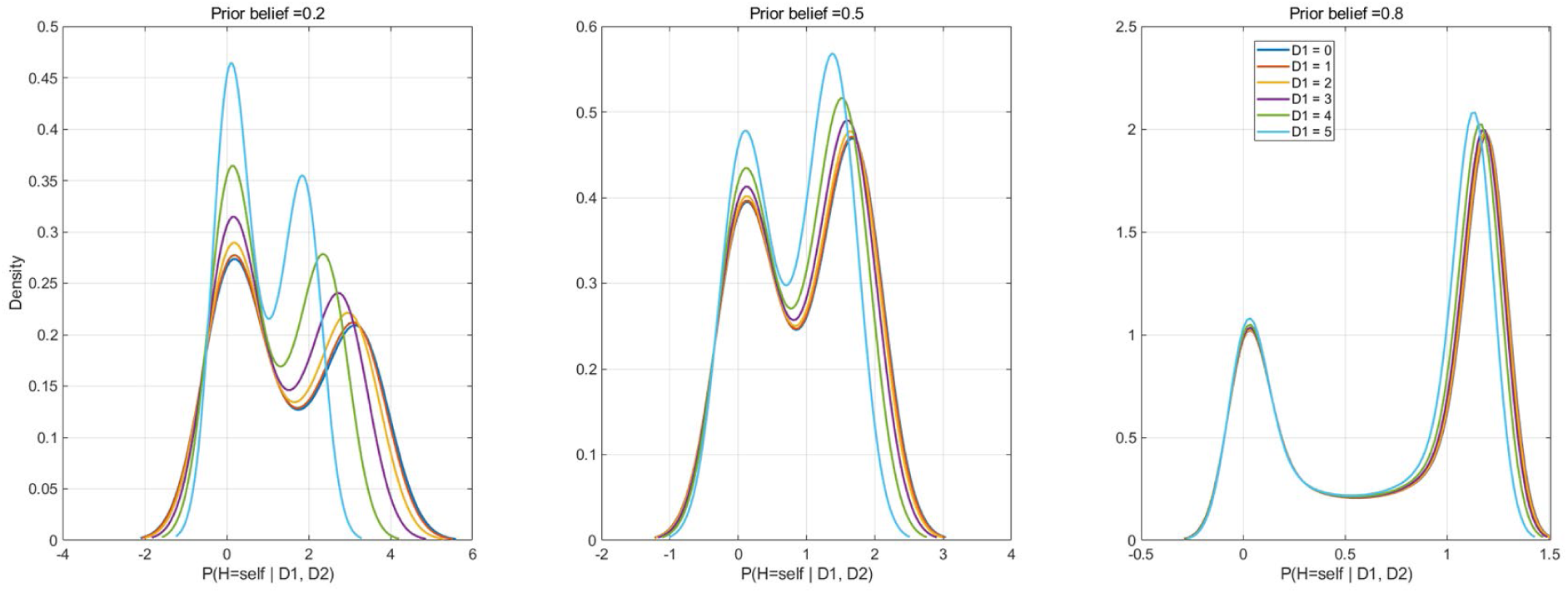
Model simulation with reduced sensorimotor sensitivity (D1) and enhanced retrospective processing (D2) while varying sensorimotor input (D1) the prior belief.

In summary, the present study proposed a Bayesian integration model that considers multiple cues and the probability of others in the environment for self-attribution. We demonstrated that the proposed model could capture the features observed in the results of a behavioral experiment, and the estimated parameters accurately reflected individual differences in the weighting of different sensory cues for self-attribution. The variance of the likelihood distribution for sensory input determines the sensitivity of the sense of agency, while the prior belief sets the criterion. We also provided an example showing that our model can effectively simulate the abnormal patterns of self-attribution observed in psychosis. The proposed model not only offers important insights into the mechanisms of the sense of agency but also serves as a powerful tool for estimating parameters that capture individual differences in self-attribution.

## Methods

### Participants

Twenty-eight and thirty university students (mean age = 22.14, range 20–26 for Experiment 1; mean age = 20.03, range 18–22 for Experiment 2) participated in Experiments 1 and 2, respectively. None of the participants took part in both experiments. All participants had normal visual acuity and motor ability. Except for two participants in Experiment 2, all were left-handed, as reported by self-assessment. The experiment was approved by local ethics committees and conducted in accordance with relevant guidelines and regulations. Written informed consent was obtained from all participants prior to the experiments, and they received financial compensation for their participation.

### Experimental tasks and procedure

Participants in Experiment 1 completed the main agency task, while participants in Experiment 2 completed both the main agency task and an additional shooting task after the main task. In the main agency task (Figure 1A), a 50-pixel dot appeared at the center of a bottom line on the screen, and a 10-degree arc (i.e., the target) moved at a speed that varied sinusoidally along a 90-degree arc trajectory. The starting position of the target was randomized for each trial. Participants were instructed to push the joystick forward to its maximum range to launch the dot, which was triggered when the joystick’s position exceeded 20%. The launching direction of the dot was influenced by both the actual joystick angle and an angular bias depending on the experimental condition (i.e., proximal error, Figure 1B). The dot reached the target’s moving trajectory in 500 ms and remained there for another 500 ms. During the dot’s movement, the program adjusted the target arc’s position, moving it toward or away from the dot’s final position according to the experimental condition (i.e., distal error, Figure 1B). Participants were informed that the target arc might occasionally change direction, and they were instructed to aim as accurately as possible at its center. A demo video showing how the dot and the target arc moved can be found at https://youtu.be/2moWdf5-fdQ.

There were five conditions for proximal error (i.e., angular bias): 0, 10, 20, 30, and 40 degrees, and five conditions for distal error (i.e., the angle between the dot’s final position and the center of the target arc, with the dot’s initial position as the origin): 0, 5, 10, 15, and 20 degrees, resulting in 25 combinations. The values of proximal and distal errors were carefully chosen based on pilot experiments to ensure participants remained motivated to aim at the target and experienced sufficient agency disruption in extreme conditions. After each trial, participants used a mouse to adjust a line scale on the screen to rate their sense of control over the dot’s trajectory (left end: no control; right end: full control). Ratings were converted to a 0–1 scale based on the position of the line.

Participants in Experiment 2 also completed an additional shooting task after the main agency task (Figure 1C). In the shooting task, the target arc remained static, and no angular bias was applied to the dot’s launching direction.

Participants were tested individually in a quiet room. They completed 10 practice trials of the agency task, followed by 150 actual trials (6 repeats for each condition), with additional practice provided upon request. The trial order was randomized. In addition, a practice shooting session, in which participants were required to aim at a static target until achieving five consecutive successes, was conducted before the main agency task practice only in Experiment 1. This likely explains why participants in Experiment 1 weighted the proximal error more than those in Experiment 2. A short break was given every 40 trials. For the shooting task in Experiment 2, no practice was conducted. The entire experiment lasted approximately 40 minutes per participant.

### Parameter Estimation for Individual Participants

We fitted the proposed model to individual behavioral results from the two experiments. For each individual, we fitted the actual values of proximal error, distal error, and agency ratings using the model to estimate the parameters σ1 and σ2 for the likelihood distributions of the two types of sensory input, as well as the prior belief in the probability of self *P*(*H*_=*self*_). Maximum likelihood estimation was applied. Parameter estimation was conducted in MATLAB R2024b with the Optimization Toolbox and Statistics and Machine Learning Toolbox (MathWorks, Inc., USA). The initial parameters for σ1, σ2, and prior belief in self were set to [20, 10, 0.5], with optimization bounds set to [1, 100] for σ and [0.1, 1] for prior belief. The code for the optimization and sample data can be found on GitLab (https://gitlab.com/wwwwen/agency-bayesian-integration).

### Measuring Sensitivity and Criterion

The sensitivity of the sense of agency to a specific sensory cue was calculated by fitting the agency ratings and the values of the sensory cues in each trial for each participant using the following logistic function. Additionally, the criterion for the sense of agency was determined by the mean agency rating for each individual. A higher agency rating indicated a more liberal tendency toward self-attribution.

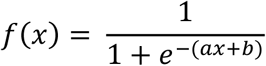

Where *x* represents the control or similarity level (e.g., the degree to which the sensory feedback aligns with self-generated action); *a* is the slope of the function, which determines the steepness of the curve; *b* shifts the curve horizontally, impacting where the inflection point (50% attribution) occurs.

## Data availability

All raw data files and experimental task code are available online at https://osf.io/4tp8x/. The code for the optimization and sample data can be found at https://gitlab.com/wwwwen/agency-bayesian-integration.

## Acknowledgements

We wish to thank Hiroyuki Hamada, Kazunori Yoshida, Yui Yamashita for their assistance with preparing the tasks and testing participants. This work was supported by JST FOREST Program (JPMJFR2144) and JST Moonshot R&D Program (JPMJMS2013).

## Author contributions

A.C-Y.C. proposed the model and analyzed all datasets. W.W. designed the experiments and collected the data with the help of people acknowledged above. A.C-Y.C. and W.W. interpreted the data and wrote the manuscript.

## References

Blakemore S-J, Wolpert DM, Frith CD. 2002. Abnormalities in the awareness of action. Trends Cogn Sci 6:237–242. doi:10.1016/S1364-6613(02)01907-1

Charalampaki A, Peters C, Maurer H, Maurer LK, Müller H, Verrel J, Filevich E. 2023. Motor outcomes congruent with intentions may sharpen metacognitive representations. Cognition 235:105388. doi:10.1016/j.cognition.2023.105388

Dakin S, Carlin P, Hemsley D. 2005. Weak suppression of visual context in chronic schizophrenia. Curr Biol 15:R822–R824. doi:10.1016/j.cub.2005.10.015

David N, Newen A, Vogeley K. 2008. The “sense of agency” and its underlying cognitive and neural mechanisms. Conscious Cogn 17:523–534. doi:10.1016/j.concog.2008.03.004

Ernst MO, Banks MS. 2002. Humans integrate visual and haptic information in a statistically optimal fashion. Nature 415:429–433. doi:10.1038/415429a

Ernst MO, Bülthoff HH. 2004. Merging the senses into a robust percept. Trends Cogn Sci 8:162–169. doi:10.1016/j.tics.2004.02.002

Frith CD, Blakemore S-J, Wolpert DM. 2000. Explaining the symptoms of schizophrenia: Abnormalities in the awareness of action. Brain Res Rev 31:357–363. doi:10.1016/S0165-0173(99)00052-1

Gallagher S. 2000. Philosophical conceptions of the self: Implications for cognitive science. Trends Cogn Sci 4:14–21. doi:10.1016/S1364-6613(99)01417-5

Haggard P. 2017. Sense of agency in the human brain. Nat Rev Neurosci 18:197–208. doi:10.1038/nrn.2017.14

Hauser M, Moore JW, de Millas W, Gallinat J, Heinz A, Haggard P, Voss M. 2011. Sense of agency is altered in patients with a putative psychotic prodrome. Schizophr Res 126:20–27. doi:10.1016/j.schres.2010.10.031

Hur J, Soo J, Young T, Park S. 2014. The crisis of minimal self-awareness in schizophrenia: A meta-analytic review. Schizophr Res 152:58–64. doi:10.1016/j.schres.2013.08.042

Koreki A, Maeda T, Fukushima H, Umeda S, Takahata K, Okimura T, Funayama M, Iwashita S, Mimura M, Kato M. 2015. Behavioral evidence of delayed prediction signals during agency attribution in patients with schizophrenia. Psychiatry Res 1–6. doi:10.1016/j.psychres.2015.08.023

Krugwasser AR, Stern Y, Faivre N, Harel EV, Salomon R. 2022. Impaired sense of agency and associated confidence in psychosis. Schizophrenia 8:32. doi:10.1038/s41537-022-00212-4

Legaspi R, Toyoizumi T. 2019. A Bayesian psychophysics model of sense of agency. Nat Commun 10:4250. doi:10.1038/s41467-019-12170-0

Leptourgos P, Corlett PR. 2020. Embodied predictions, agency, and psychosis. Front Big Data 3:27. doi:10.3389/fdata.2020.00027

Leube DT, Knoblich G, Erb M, Schlotterbeck P, Kircher TTJJ. 2010. The neural basis of disturbed efference copy mechanism in patients with schizophrenia. Cogn Neurosci 1:111–117. doi:10.1080/17588921003646156

Lindner A, Thier P, Kircher TTJ, Haarmeier T, Leube DT. 2005. Disorders of agency in schizophrenia correlate with an inability to compensate for the sensory consequences of actions. Curr Biol 15:1119–24. doi:10.1016/j.cub.2005.05.049

Mariano M, Rossetti I, Maravita A, Paulesu E, Zapparoli L. 2024. Sensory Attenuation Deficit and Auditory Hallucinations in Schizophrenia: A Causal Mechanism or a Risk Factor? Evidence From Meta-Analyses on the N1 Event-Related Potential Component. Biol Psychiatry 96:207–221. doi:10.1016/j.biopsych.2023.12.026

McGuire PK, Silbersweig DA, Wright I, Murray RM, David AS, Frackowiak RSJ, Frith CD. 1995. Abnormal monitoring of inner speech: A physiological basis for auditory hallucinations. Lancet 356:596–600.

Metcalfe J, Eich TS, Miele DB. 2013. Metacognition of agency: Proximal action and distal outcome. Exp Brain Res 229:485–496. doi:10.1007/s00221-012-3371-6

Metcalfe J, Van Snellenberg JX, DeRosse P, Balsam P, Malhotra a. K. 2012. Judgements of agency in schizophrenia: An impairment in autonoetic metacognition. Philos Trans R Soc B Biol Sci 367:1391–1400. doi:10.1098/rstb.2012.0006

Moore JW. 2016. What is the sense of agency and why does it matter. Front Psychol 00:00. doi:10.3389/fpsyg.2016.01272

Moore JW, Fletcher PC. 2012. Sense of agency in health and disease: A review of cue integration approaches. Conscious Cogn 21:59–68. doi:10.1016/j.concog.2011.08.010

Oi H, Wen W, Chang AY-C, Uchida H, Maeda T. 2024. Hierarchical analysis of the sense of agency in schizophrenia: motor control, control detection, and self-attribution. Schizophrenia 10:79. doi:10.1038/s41537-024-00512-x

Pacherie E. 2008. The phenomenology of action: A conceptual framework. Cognition 107:179–217. doi:10.1016/j.cognition.2007.09.003

Posada A, Franck N, Augier S, Georgieff N, Jeannerod M. 2007. Altered processing of sensorimotor feedback in schizophrenia. Comptes Rendus - Biol 330:382–388. doi:10.1016/j.crvi.2007.02.003

Powers AR, Kelley M, Corlett PR. 2016. Hallucinations as top-down effects on perception. Biol Psychiatry Cogn Neurosci Neuroimaging 1:393–400. doi:10.1016/j.bpsc.2016.04.003

Renes RA, Vermeulen L, Kahn RS, Aarts H, van Haren NEM. 2013. Abnormalities in the establishment of feeling of self-agency in schizophrenia. Schizophr Res 143:50–54. doi:10.1016/j.schres.2012.10.024

Rollwage M, Dolan RJ, Fleming SM. 2018. Metacognitive failure as a feature of those holding radical beliefs. Curr Biol 28:4014–4021.e8. doi:10.1016/j.cub.2018.10.053

Rossetti I, Mariano M, Maravita A, Paulesu E, Zapparoli L. 2024. Sense of agency in schizophrenia: A reconciliation of conflicting findings through a theory-driven literature review. Neurosci Biobehav Rev 163:105781. doi:10.1016/j.neubiorev.2024.105781

Roth MJ, Lindner A, Hesse K, Wildgruber D, Wong HY, Buehner MJ. 2023. Impaired perception of temporal contiguity between action and effect is associated with disorders of agency in schizophrenia. Proc Natl Acad Sci 120. doi:10.1073/pnas.2214327120

Rubio JL, Ruiz-Veguilla M, Hernández L, Barrigón ML, Salcedo MD, Moreno JM, Gómez E, Moritz S, Ferrín M. 2011. Jumping to conclusions in psychosis: A faulty appraisal. Schizophr Res 133:199–204. doi:10.1016/j.schres.2011.08.008

Schneider K. 1959. Clinical Psychopathology. New York: Grune and Stratton.

Stanghellini G, Monti MR. 1993. Influencing and being influenced: The other side of ‘bizarre delusions’. 2: Clinical investigation. Psychopathology 26:165–169.

Synofzik M, Vosgerau G, Lindner A. 2009. Me or not me - An optimal integration of agency cues? Conscious Cogn 18:1065–1068. doi:10.1016/j.concog.2009.07.007

Tan DPW, Carter O, Marshall D, Perrykkad K. 2023. Agency in schizophrenia and autism: a systematic review. Front Psychol 14:1280622. doi:10.3389/fpsyg.2023.1280622

Uhlmann L, Pazen M, van Kemenade BM, Kircher T, Straube B. 2021. Neural correlates of self-other distinction in patients with schizophrenia spectrum disorders: The roles of agency and hand identity. Schizophr Bull 47:1399–1408. doi:10.1093/schbul/sbaa186

Vallacher RR, Wegner DM. 1989. Levels of Personal Agency: Individual Variation in Action Identification. J Pers Soc Psychol 57:660–671. doi:10.1037/0022-3514.57.4.660

Vinding MC, Pedersen MN, Overgaard M. 2013. Unravelling intention: Distal intentions increase the subjective sense of agency. Conscious Cogn 22:810–815. doi:10.1016/j.concog.2013.05.003

Voss M, Moore JW, Hauser M, Gallinat J, Heinz A, Haggard P. 2010. Altered awareness of action in schizophrenia: A specific deficit in predicting action consequences. Brain 133:3104–3112. doi:10.1093/brain/awq152

Wegner DM. 2003. The mind’s best trick: How we experience conscious will. Trends Cogn Sci 7:65–69. doi:10.1016/S1364-6613(03)00002-0

Wegner DM, Sparrow B, Winerman L. 2004. Vicarious agency: Experiencing control over the movements of others. J Pers Soc Psychol 86:838–848. doi:10.1037/0022-3514.86.6.838

Wen W. 2019. Does delay in feedback diminish sense of agencyA review. Conscious Cogn 73:102759. doi:10.1016/j.concog.2019.05.007

Wen W, Chang AY-C, Imamizu H. 2024. The sensitivity and criterion of sense of agency. Trends Cogn Sci 28:397–399. doi:10.1016/j.tics.2024.03.002

Wen W, Imamizu H. 2022. The sense of agency in perception, behaviour and human–machine interactions. Nat Rev Psychol 1:211–222. doi:10.1038/s44159-022-00030-6

Wen W, Ishii H, Ohata R, Yamashita A, Asama H, Imamizu H. 2021. Perception and control: individual difference in the sense of agency is associated with learnability in sensorimotor adaptation. Sci Rep 11:1–8. doi:10.1038/s41598-021-99969-4

Wen W, Kuroki Y, Asama H. 2019. The sense of agency in driving automation. Front Psychol 10:02691. doi:10.3389/fpsyg.2019.02691

Wen W, Yamashita A, Asama H. 2015. The influence of goals on sense of control. Conscious Cogn 37:83–90. doi:doi:10.1016/j.concog.2015.08.012

Wolpert DM, Flanagan JR. 2001. Motor prediction. Curr Biol 11:R729–R732. doi:10.1016/S0960-9822(01)00432-8

Wright JL, Chen JYC, Barnes MJ. 2018. Human–automation interaction for multiple robot control: the effect of varying automation assistance and individual differences on operator performance. Ergonomics 61:1033–1045. doi:10.1080/00140139.2018.1441449

