## Supplementary Material S1 for "Bayesian Integration in Sense of Agency: Understanding Self-attribution and Individual Differences"

**S1**. Probability density distribution of agency judgment for each participant (n = 58) across all trials from the two experiments when varying the proximal error (A) and the distal error (B). The results revealed large individual differences in both the weightings of each type of cue and the boundary of sense of agency. To enhance visualization of these distributions, kernel density estimation was applied.

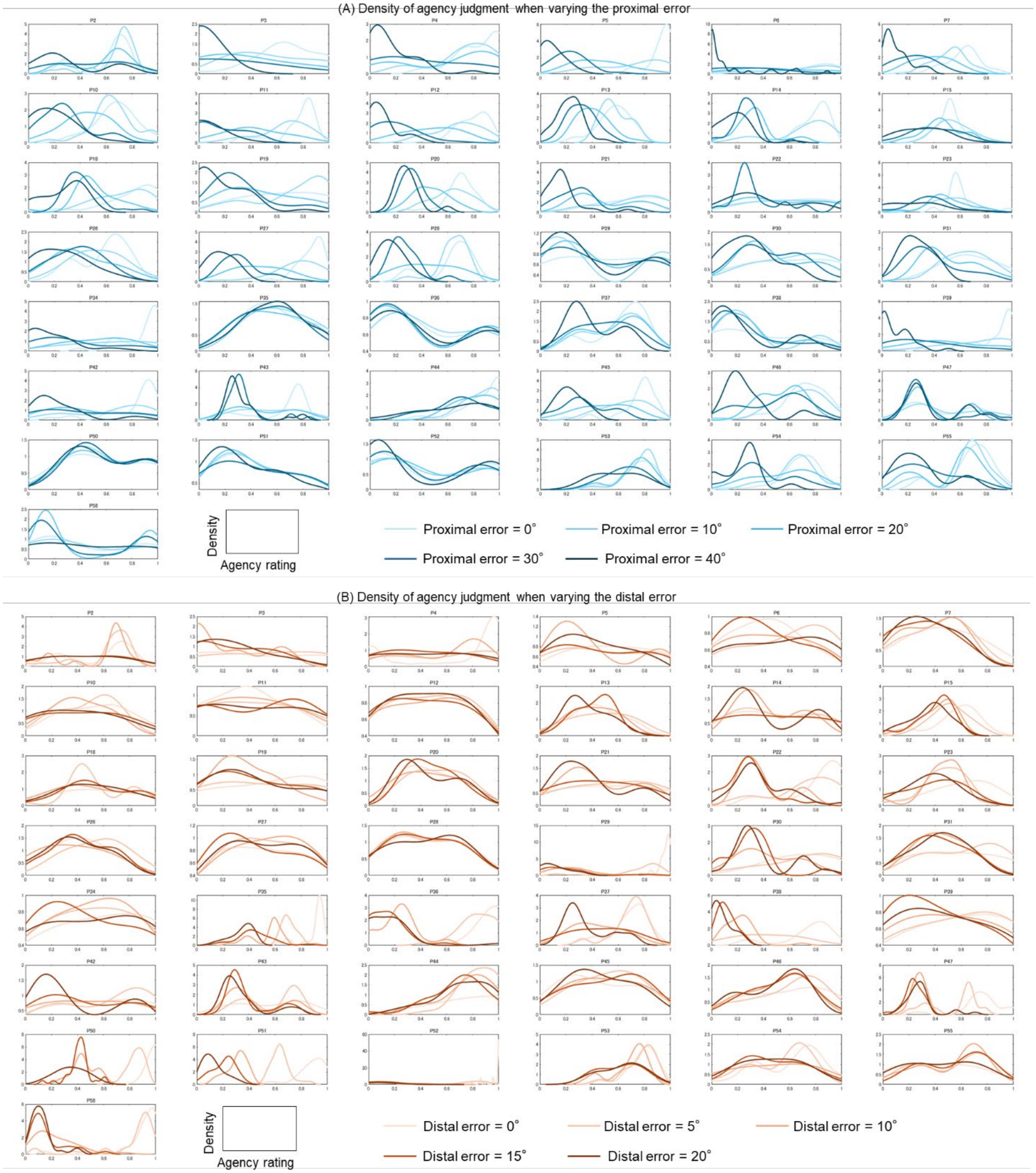
